# OneFlowTraX: A User-Friendly Software for Super-Resolution Analysis of Single-Molecule Dynamics and Nanoscale Organization

**DOI:** 10.1101/2023.08.10.552827

**Authors:** Leander Rohr, Alexandra Ehinger, Luiselotte Rausch, Nina Glöckner Burmeister, Alfred J. Meixner, Julien Gronnier, Klaus Harter, Birgit Kemmerling, Sven zur Oven-Krockhaus

## Abstract

Super-resolution microscopy (SRM) approaches revolutionize cell biology by providing insights into the nanoscale organization and dynamics of macromolecular assemblies and single molecules in living cells. A major hurdle limiting SRM democratization is post-acquisition data analysis which is often complex and time-consuming. Here, we present OneFlowTraX, a user-friendly and open-source software dedicated to the analysis of single-molecule localization microscopy (SMLM) approaches such as single-particle tracking photoactivated localization microscopy (sptPALM). Through an intuitive graphical user interface, OneFlowTraX provides an automated all-in-one solution for single-molecule localization, tracking, as well as mobility and clustering analyses. OneFlowTraX allows the extraction of diffusion and clustering parameters of millions of molecules in a few minutes. Finally, OneFlowTraX greatly simplifies data management following the FAIR (Findable, Accessible, Interoperable, Reusable) principles. We provide a detailed step-by-step manual and guidelines to assess the quality of single-molecule analyses. Applying different fluorophores including mEos3.2, PA-GFP, and PATagRFP, we exemplarily used OneFlowTraX to analyze the dynamics of plant plasma membrane-localized proteins including an aquaporin, the brassinosteroid receptor Brassinosteroid Insensitive 1 (BRI1) and the Receptor-Like Protein 44 (RLP44).

## INTRODUCTION

All living cells regulate the dynamics and organization of molecules at the nanoscale to control their biological processes. Accordingly, appropriate methods are needed to elucidate the underlying mechanisms and functions at the molecular level. In recent years, there has been a significant focus on the plasma membrane (PM) due to its crucial role in important functions such as homeostasis and mass transport, and its primary role as a mediator of signals into and out of the cell. However, only a few techniques allow the in vivo analysis of molecules with high spatiotemporal resolution. Suitable microscopy techniques include fluorescence recovery after bleaching (FRAP) (Axelrod et al., 1976; Peters et al., 1974), fluorescence correlation spectroscopy (FCS) (Magde et al., 1972), and single-particle tracking (spt) (Manzo and Garcia-Parajo, 2015) of labeled molecules, the latter being mainly driven by the advent of super-resolution technologies such as photoactivated localization microscopy (PALM) (Betzig et al., 2006; Hess et al., 2006; Manley et al., 2008).

Imaging of single molecules in living cells is usually performed under total internal reflection (TIRF) illumination, which provides greatly enhanced contrast for a thin layer of the biological sample due to its small penetration depth of around 150 nm beyond the coverslip. However, due to the relatively thick cell wall of plant cells, their compartments, such as the PM, are not in the optimal range for TIRF. Therefore, alternative illumination methods such as highly inclined thin illumination (HiLo) (Tokunaga et al., 2008), also known as variable angle epifluorescence microscopy (VAEM) (Konopka and Bednarek, 2008), are widely used for plants. Moreover, due to the limited permeability of the cell wall, plant cell biologists cannot use organic dyes common in the animal or human field (Lelek et al., 2022) for live cell imaging but must rely on a limited selection of genetically encoded fluorophores fused to the gene of interest (Hosy et al., 2015; Jolivet et al., 2023; McKenna et al., 2019).

These technical difficulties have contributed to the fact that dynamic analysis of proteins in plant cells at very high resolution has only recently taken off. The improvement of technical possibilities in microscopy and other methods, such as single-molecule tracking and cluster analysis, now offers data on dynamic parameters, including diffusion coefficients and nanoscale organization, especially for PM proteins (Bayle et al., 2021; Gronnier et al., 2017; Gronnier et al., 2019; Hosy et al, 2015; Jaillais and Ott, 2020; Martiniere and Zelazny, 2021; McKenna et al, 2019; Perraki et al., 2018; Smokvarska et al., 2023).

Despite the progress made in recent years, the analysis of single-molecule localization microscopy (SMLM) data remains a complicated, multistep process. First, the positions of the individual labeled membrane proteins in each image must be determined with high precision (localization), followed by the assignment of localizations across multiple images to trajectories (tracking). The mobilities or proportions of mobile/immobile proteins are then calculated from the analysis of these trajectories. Subsequently, a map of all observed single molecules localizations can be reconstituted to analyze the nanoscale organization of molecules (e.g., cluster analysis). Each of these steps has been implemented over the years with dedicated analysis software (Chenouard et al., 2014; Sage et al., 2019). Localization depends on physical camera parameters and localization algorithms that must incorporate noise statistics for the low photon counts typical of single-molecule microscopy. Tracking can be performed with a variety of algorithms, and the chosen parameters have a large impact on the data evaluation, representation and thus their interpretation. Mobility analysis is initially performed using mean square displacement (MSD) plots (Qian et al., 1991). Still, this method is not always applicable and is highly sensitive to the parameters used. Combining sptPALM data with cluster analysis is a relatively recent development; several methods have advantages and disadvantages, leading to different data evaluation, representation and interpretation. This procedural complexity and challenges prevent broader application of SMLM techniques such as sptPALM especially in plant cell biology.

Although practical guides have been published recently (Bayle et al, 2021), there is still a lack of software packages that guide the user through all analyses without excessive computational knowledge. The available software is usually limited to or specializes in only one of the above sub-steps, with compatibility problems between the different solutions.

In this work we present a comprehensive, all-in-one open-source, time-saving software package, named OneFlowTraX, guiding the scientist through the steps of SMLM-based localization, tracking, mobility- and cluster analyses of molecules in living cells, which we apply to plant cells. Moreover, the storage of the SMLM data and metadata follow FAIR (Findable, Accessible, Interoperable and Reusable) principles (Wilkinson et al., 2016) for scientific data management and stewardship. We explored three genetically encoded fluorophores (mEos3.2, PA-GFP and PATagRFP) of different photophysical characteristics, suitable for sptPALM studies in living plant cells, exemplified by the temporal and spatial analysis of four different plant plasma membrane proteins. The PA-GFP and PATagRFP pair will also enable dual-color sptPALM applications in the future.

## METHOD

### Design and properties of OneFlowTraX

#### Purpose and workflow

OneFlowTraX runs as an executable program or as an application in MATLAB (Mathworks). The user is guided through several steps, including the localization of single molecules, the reconstruction of single-molecule trajectories, the calculation of mobility parameters, and cluster analyses based on molecule or trajectory positions. The easily scriptable and customizable code is available on GitHub (https://github.com/svenzok/OneFlowTraX), accompanied by a user guide that answers all analysis-specific questions. For each main analysis step (localization, tracking, mobility and cluster analysis), extensive literature research was conducted to select the most suitable method or algorithm for OneFlowTraX. Published codes were partially adapted to meet the requirements of OneFlowTraX; alternatively, new software code was written to provide a robust and efficient all-in-one analysis pipeline for state-of-the-art single-molecule imaging analyses. This eliminates the need for the user to convert analysis data from one specialized software solution to another, which both saves time and reduces the complexity of these consecutive analysis steps. Some combinations, such as the generation and use of track data for cluster analysis (see below), are particularly important for the evaluation of single-molecule tracking experiments and were not previously available in an integrated software pipeline. The individual steps are discussed in more detail in the following sections.

#### Localization

The sptPALM raw data (Figure 1A) usually consists of time series images that are saved as TIFF image stacks. Typical formats in our experiments are 100×100 to 400×400 pixels (corresponding to 10×10 to 40×40 µm) with 2 000 to 10 000 images taken at a frame rate between 20 and 50 Hz. These parameters can vary depending on the specific experiment but are mainly governed by the optimal magnification of the imaging system for single-molecule detection, the choice of the frame rate that achieves sufficient contrast, the size of the imaged PM region of interest, and the photophysical characteristics of the chosen fluorophore.

**Figure 1.**
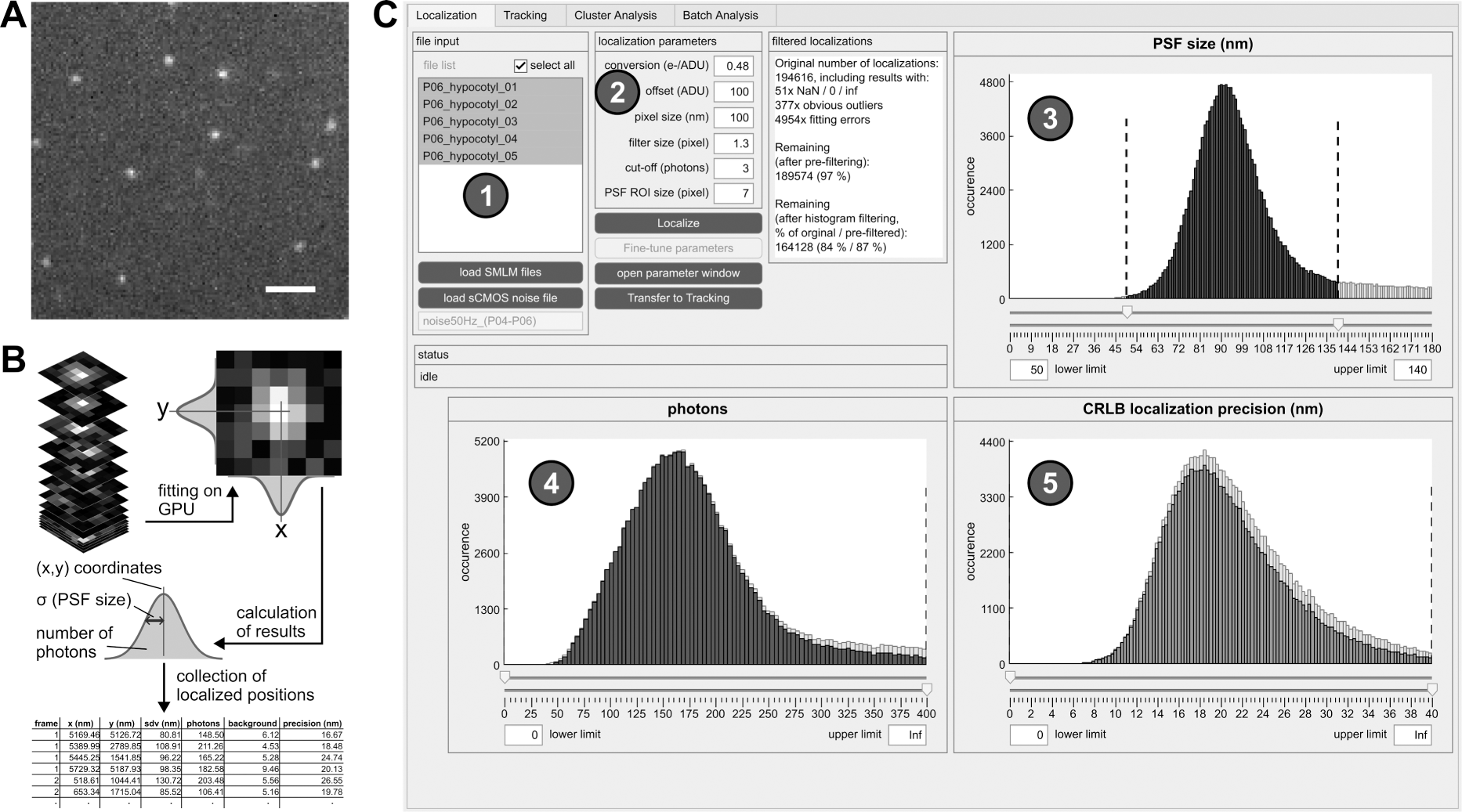
Single-molecule localization procedure. **(A)** Single frame of a typical sptPALM image stack. Scale bar = 2 µm. **(B)** Localization process. Each putative single molecule position is cut out as a larger section and fitted with a two-dimensional Gaussian function. The fitting results (most importantly, the coordinates) for each detected single molecule are stored as lists. **(C)** Graphical user interface (GUI) of OneFlowTraX for the localization step. After loading in the sptPALM files (1), the localization procedure is executed based on user-defined parameters (2), with its results visualized as three histograms (PSF sizes (3), photon numbers (4) and localization precision (5)) for all detected and fitted fluorescence spots. This serves as quality control and allows the user to filter out, e.g., badly localized single-molecule positions.

In the first step, the fluorophore-tagged molecules are localized in each image with high precision using algorithms developed for super-resolution microscopy. Bright pixels indicate possible detected single molecules, and larger sections around these pixels are fitted with a two-dimensional Gaussian (Figure 1A). This results not only in the position but also the PSF size, the number of photons and the subsequently calculated localization accuracy, which are stored for each localized single molecule (Figure 1B). For this purpose, we adapted the core of the SMAP software (Ries, 2020). Its algorithm for determining single-molecule positions uses a robust maximum likelihood estimation (MLE) of Gaussian point spread function (PSF) models, which was shown to be very accurate (Sage et al, 2019). Furthermore, it can make use of a graphic processing unit (GPU) that massively accelerates image processing speed. In addition, the fitter can also account for the pixel-specific noise of commonly used complementary metal-oxide semiconductor (CMOS) cameras. In OneFlowTraX, this process can be started for a list of files using default parameters (Figure 1C). An auxiliary window can be opened to check the performance of the algorithm on selected single images and adjust the parameters (see the user guide that is provided with the software for detailed information) if necessary. At the end of the localization process, three histograms show (i) the PSF sizes, (ii) photons and (iii) localization accuracy for all detected and fitted molecules. These histograms can be used to check the quality of the raw data and exclude outliers from further analysis, for example, localizations with unusually large PSFs due to poor focusing or aggregation artifacts. After review, the molecule positions can be used to define single-molecule trajectories.

#### Tracking

The generally low localization density in sptPALM data allows for a simple but very fast tracking algorithm. A trajectory is formed from all those molecule positions that do not exceed a maximum distance from each other in successive images (Figure 2A). In addition, short sequences are bridged in which a molecule was temporarily undetectable (gap closing), which is typical of the blinking behavior of fluorophores observed in single-molecule microscopy. The corresponding code for this algorithm was adapted from a MATLAB program published by Jean-Yves Tinevez (Tinevez, 2011). The underlying Linear Assignment Problem (LAP) tracker (Jaqaman et al., 2008) is also part of the widely used software TrackMate (Tinevez et al., 2017). After performing the tracking in OneFlowTraX, the resulting trajectories can be visualized (Figure 2B, upper right), and colored according to their duration, displacement, mobility and other characteristics. In case of obvious connection errors (see the user guide for examples), the tracking can be repeated with adjusted track-building parameters.

**Figure 2.**
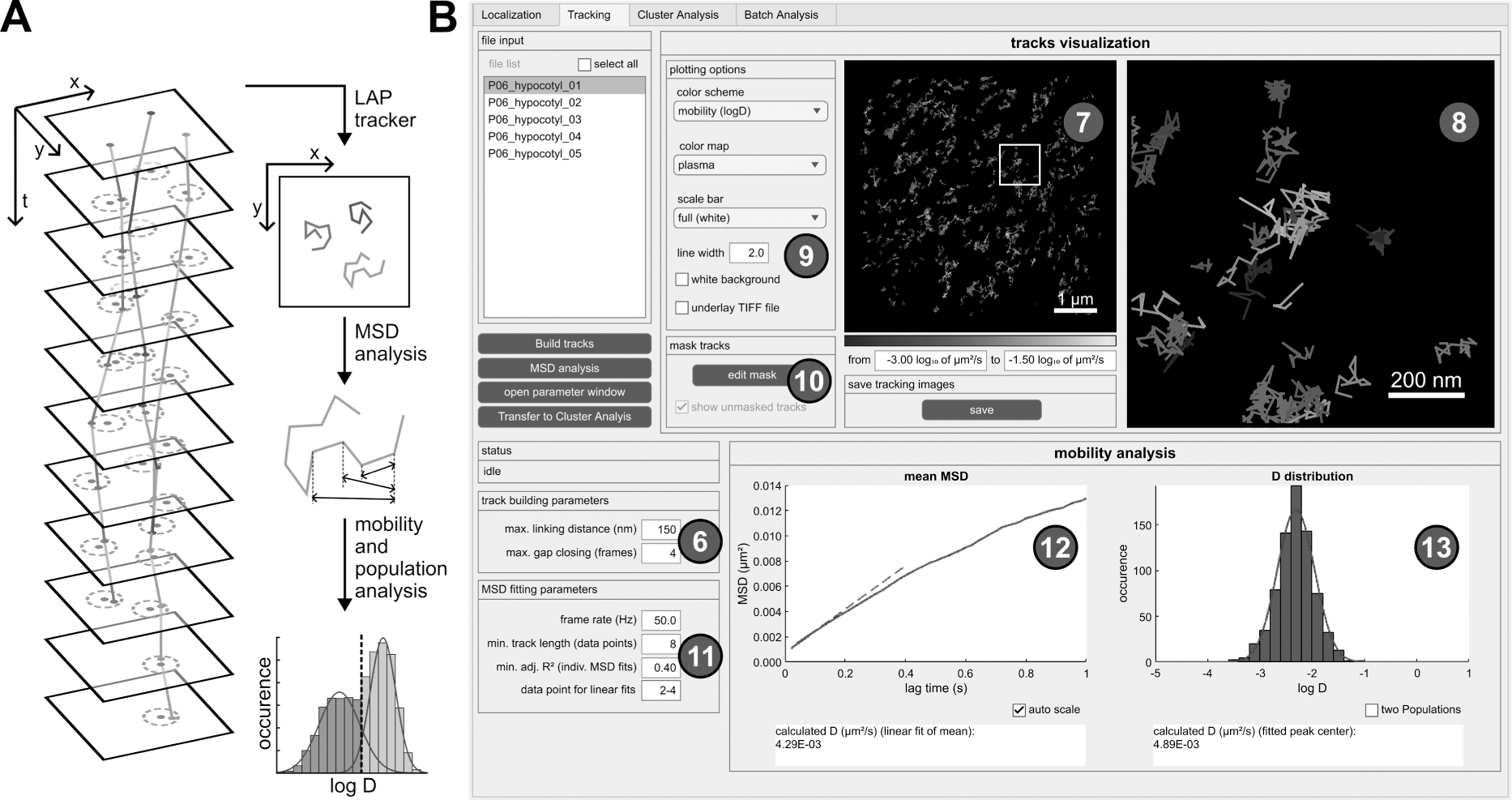
Tracking procedure and mobility analysis. **(A)** Schematic overview of the tracking procedure and subsequent analyses. Single-molecule positions are connected to trajectories when they do not exceed a maximum distance (dashed circles) in subsequent frames. The Linear Assignment Problem (LAP) tracker uses a global optimization and can also deal with detection gaps. All tracks then undergo mean square displacement (MSD) analysis based on averaged traveled distances for certain time intervals, offering mobility metrics like diffusion coefficients and more extensive evaluations such as population analyses. **(B)** GUI of OneFlowTraX for the tracking step, including mobility analysis. The track building algorithm uses the specified parameters (6) to connect the detected localization to trajectories, which are displayed in the overview image (7) and a user-defined magnified view (8). Different layers and visualization options can be chosen (9), and the resulting images stored as vector graphics. A binary mask can be defined for each file (10) to exclude tracks (for example, extracellular areas) from further analysis. The mobility analysis via MSD plots is performed based on user-specified parameters (11) and visualized as the mean MSD plot (12), containing the averaged MSD data of the selected file(s). Additionally, the distribution of logarithmized, individual diffusion coefficients (13) is shown, based on the collection of derived diffusion coefficients from fitting the MSD curves of each individual track.

#### Mobility analysis

The mean square displacement (MSD) analysis is currently the most widely used method for extracting diffusion coefficients and motion patterns for this type of data (Manzo and Garcia-Parajo, 2015). Therefore, the distances a molecule has traveled in certain time intervals is assessed to calculate mobility metrics like the diffusion coefficient (Figure 2A). A more detailed description of this method was described by Saxton and Jacobson, 1997. Only selected tracks with a certain minimum length (typically eight or more localized positions) are used for further analyses, removing short track artifacts that may originate from background signals. The mean MSD over all tracks is calculated and the diffusion coefficient is estimated via a linear fit (Figure 2B, lower left plot) that commonly only includes the first few points of an MSD curve. While this averaged analysis gives a first impression of protein mobility, a more detailed analysis is possible by estimating individual diffusion coefficients for the MSD curves of each track. Their distribution can then be plotted as a histogram (Figure 2B, lower right plot), revealing the potential existence of nonuniform mobility distributions that would remain hidden in the mean MSD plot. Due to the small number of data points for individual tracks, a goodness-of-fit threshold value (the adjusted R²) can be specified for the linear fit to reject inconclusive results. In addition to storing the peak log(D) value for each file examined, the relative proportion of multiple populations can also be estimated. Performing the MSD analysis adds new options to the tracking images, such as coloring tracks based on their mobility or splitting them visually into mobile and immobile tracks (for more details refer to the user guide).

#### Cluster analysis

While mobility analysis provides information about the dynamics of membrane proteins, their spatial nanoscale organization can also be obtained by cluster analysis based on the single-molecule data using OneFlowTraX, which implements several current state-of-the-art methods. Our analysis pipeline includes Voronoi tessellation (Andronov et al., 2016; Levet et al., 2015), density-based spatial clustering of applications with noise (DBSCAN) (Ester et al., 1996) and the recently introduced nanoscale spatiotemporal indexing clustering (NASTIC) (Wallis et al., 2023) (Figure 3A). A detailed description of these methods can also be found in the user guide.

**Figure 3.**
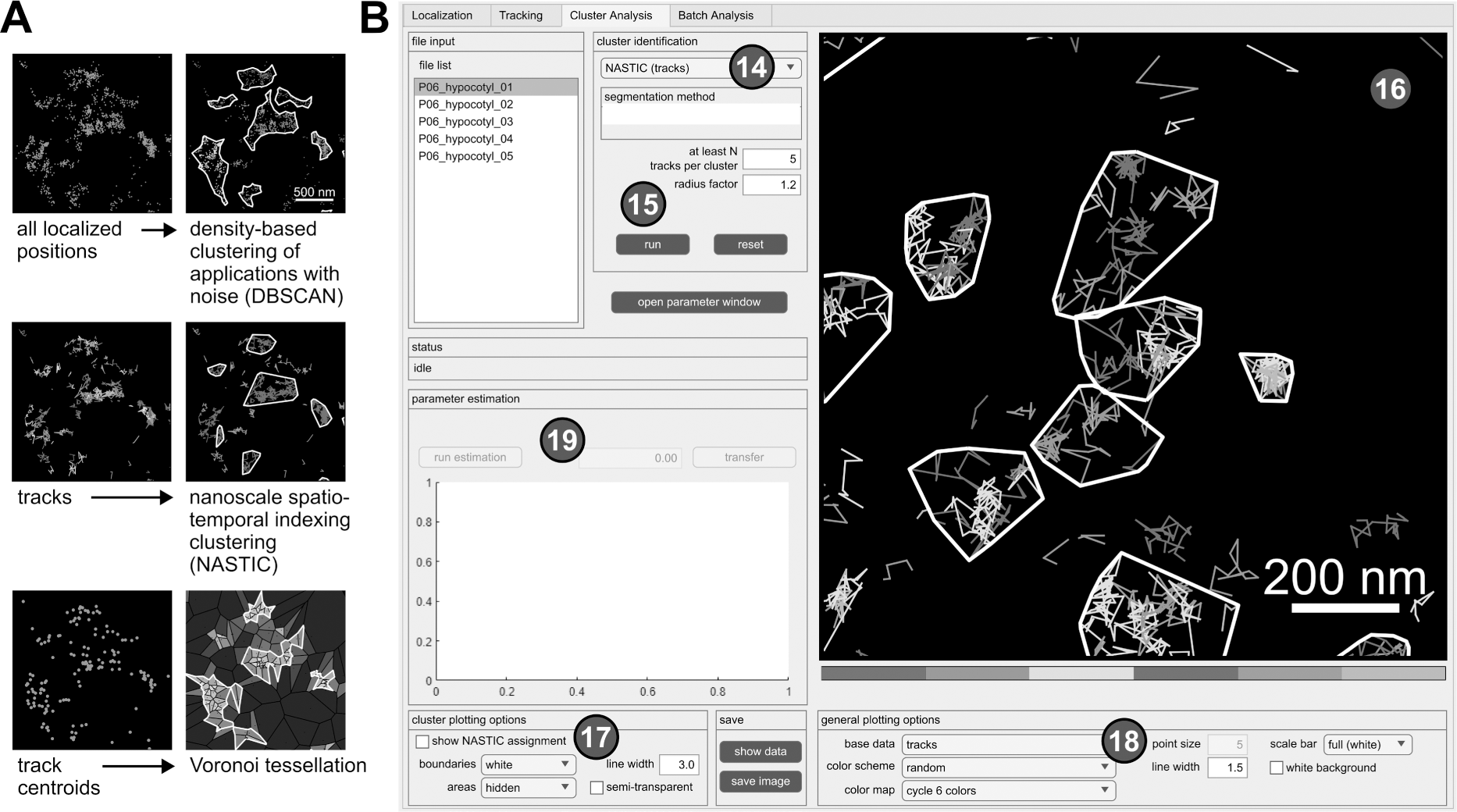
Application of clustering algorithms. **(A)** Different representations of the same exemplary data set (all localized positions, tracks and track centroids), and visualized results of implemented clustering algorithms. **(B)** GUI of OneFlowTraX for the cluster analysis. The user can choose from several clustering algorithms (14) and set their specific parameters (15). The visualization of the results (16) depends on the chosen method and can be adjusted with several options (17), (18). For some algorithms, parameter estimates can be calculated (19) based on the chosen data.

When applying the different clustering methods, the user can base the analysis in OneFlowTraX (Figure 3B) on two different data sets: (i) all localized fluorophore positions in the sample or (ii) the centroids of the assembled tracks. Option (i) disregards the assignment of localizations to individual membrane proteins, which can result in varying numbers of cluster points for each protein, while option (ii) assigns each detected membrane protein to one cluster point, at the expense of a smaller set of points (also compare Figure 3A). This choice largely depends on the available data quantity, but option (ii) should be generally preferred for its more consistent assignment. Only the NASTIC method is based solely on the protein tracks themselves, since their spatial overlap is used for cluster analysis. OneFlowTraX implements all the above algorithms, so the users can follow their preferences or compare different algorithms. This use of calculated track data for cluster analysis in a single pipeline has, to our knowledge, not been available before. It can therefore greatly benefit the data analysis of single particle tracking experiments, not only in terms of processing speed, but also by providing control and knowledge of upstream analysis step parameters that could influence the cluster analysis results. Finally, all applied analyses are visualized and can be color-coded with different settings and exported.

#### Batch processing

The batch analysis, outlined schematically in Figure 4A, automatically processes the raw sptPALM data selected by the user and then stores all analysis results in a consistent format. It allows the evaluation of big datasets, including all the above analysis steps, in a very short time (about seven seconds per file with approximately 30 000 localizations each). A summary of all parameters used for the individual steps in the analysis of the membrane protein of interest is listed in an intuitive selection structure (Figure 4B). If necessary, parameters can still be changed here, and the subsequent batch analysis is performed based on these parameters collectively for all analysis steps (localization, tracking, mobility and cluster analysis). The parameter list is also attached to the collected results so that all steps and settings can be traced back according to the FAIR principle.

**Figure 4.**
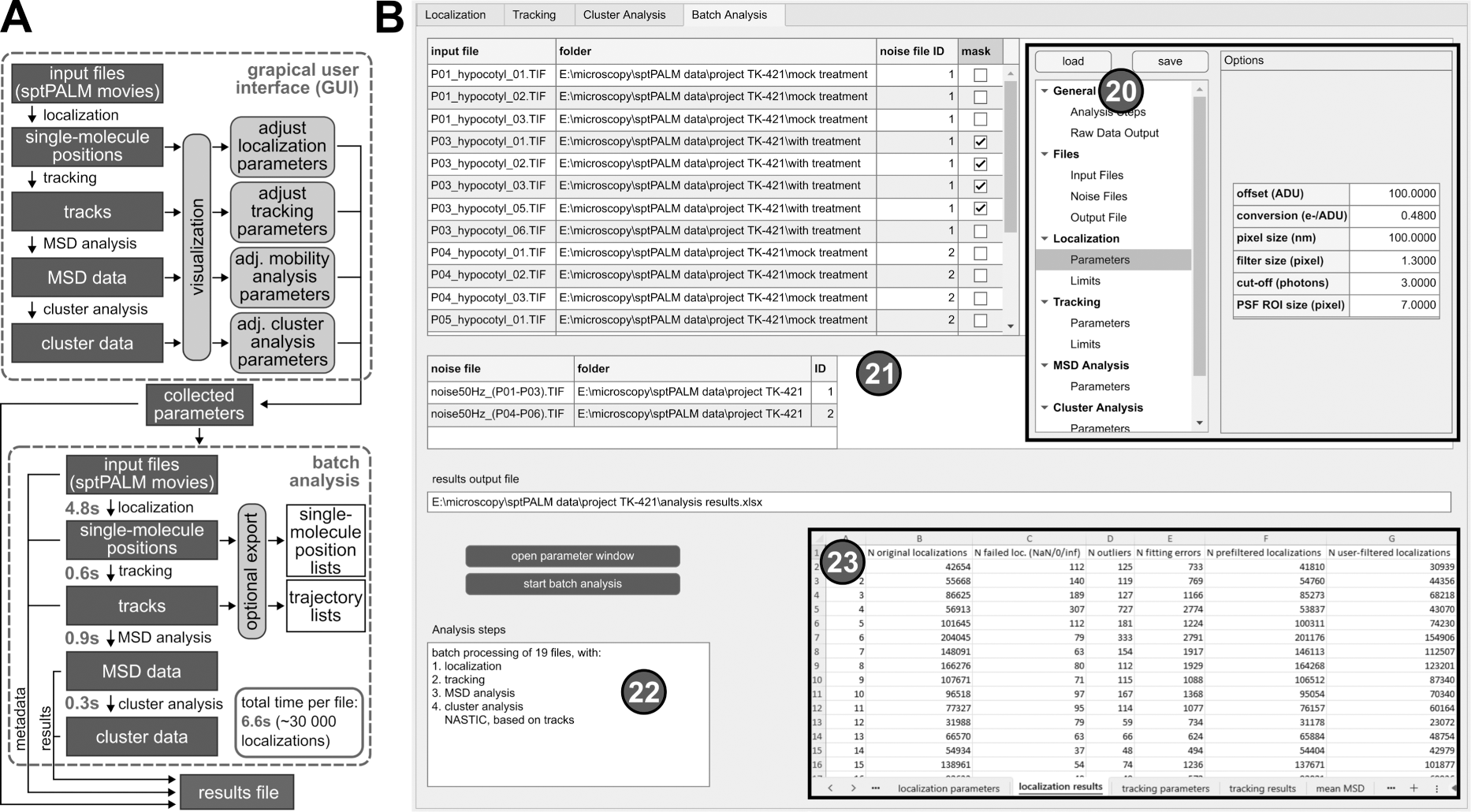
Workflow and batch processing. **(A)** Schematic overview of the workflow steps in the GUI of the software and the batch analysis. The GUI is intended to find a suitable set of parameters for all analysis steps, supported by the visualization of intermediate results. The collected parameters can then be used in the batch analysis, which quickly performs all analysis steps for each input file and saves all relevant data in one comprehensive results file. The measured fitting times per file were achieved using an AMD Ryzen 5 4500 CPU and a GTX 960 consumer graphics card **(B)** GUI of OneFlowTraX for the batch analysis. All user-specified parameters from all previous tabs are stored in a systematic overview window (20) that can be opened from any tab. Input files for batch processing are also added here and are shown as tables (21) for inspection and the assignment of noise files and masks. The entire file list can then be automatically processed, beginning with the localization procedure and may include tracking, mobility and cluster analysis as chosen by the user, summarized in (22). All parameters and results are stored in a spreadsheet file, a section of which is shown as the inset (23).

### Fluorophores for in planta sptPALM analysis

In plant cells, the rather impenetrable cell wall largely precludes dye- or quantum dot based single molecule tracking approaches, so that sptPALM with genetically encoded fluorescent proteins (FPs) must be used, which are fused translationally to the protein of interest. Conversely, FPs exhibit comparatively low photostability and brightness. Moreover, sptPALM analyses of proteins require photoactivatable or -convertible FP variants, where the density of visible fluorophores per image is controlled by an additional activation laser, further limiting the pool of applicable FPs. Genetic constructs encoding photoconvertible mEos2 fusions have been useful for sptPALM-based analyses of membrane proteins in plant cells (Gronnier et al, 2017; Hosy et al, 2015; Smokvarska et al, 2023). However, it was shown previously that mEos2 tends to form oligomers and aggregates in animal cells, especially when fused to membrane proteins (Zhang et al., 2012). We therefore recommend to prefer the improved version mEos3.2 that is monomeric and also works in plants (Jolivet et al, 2023). As with mEos2, the native green form of mEos3.2 can be converted to a red form using light of ∼400 nm wavelength, allowing to adjust the density of the visible fluorophores in the red imaging channel. However, as mEos3.2 occupies both the green and red parts of the spectrum, it is not compatible to combine it with other FPs for potential dual-color applications. Based on their photophysical characteristics (Supplemental Table 1), we propose the use of photoactivatable (PA-) GFP (Patterson and Lippincott-Schwartz, 2002) and PATagRFP (Subach et al., 2010) as additional fluorophores for sptPALM applications in plant cells. Both PA FPs are non-fluorescent in their respective native forms and can be converted to their spectrally distinct fluorescent forms with light of ∼400 nm, enabling simultaneous imaging of differently labeled proteins in two color channels. The three fluorophore coding sequences were codon-optimized for their use in plant cells (Supplemental Table 2).

## RESULTS

### PM proteins used for proof-of-principle sptPALM analyses

In order to demonstrate the applicability of OneFlowTraX for sptPALM analyses with focus on plant cells, well-described *A. thaliana* membrane proteins were used, namely BRI1, RLP44, LTi6a and the aquaporin PIP2;1. LTi6a-mEos2 and PIP2;1-mEos2 *A. thaliana* transgenic lines (Hosy et al, 2015) are under the control of the *PIP2;1* promoter (*pPIP2;1*). BRI1 and RLP44 were expressed as mEos3.2, PA-GFP and PATagRFP fusions under the control of the respective native promoter (*pBRI1*, *pRLP44*).

The aquaporin PIP2;1 is a large six transmembrane domains-containing, tetrameric water and hydrogen peroxide permeable pore (Dynowski et al., 2008), whereas LTi6a is a small two transmembrane domains-containing intrinsic PM protein of yet unclear function (Kim et al., 2021). BRI1 initiates well-understood signaling pathways in plant cells. Upon binding of BR to BRI1’s extracellular domain, its interaction with the co-receptor BRI1-ASSOCIATED KINASE 1 (BAK1) is enhanced. This leads to a re-arrangement of proteins within the complex, eventually resulting in auto- and transphosphorylation of their kinase domains and its full signaling activity (Gou and Li, 2020; Wolf, 2020). The outcomes of BR activation of the BRI1/BAK1 complex are on the one hand the differential regulation of BR-responsive genes via a nucleo-cytoplasmic signaling cascade (Mora-Garcia et al., 2004; Vert and Chory, 2006; Yin et al., 2005; Zhu et al., 2017) and on the other hand a rapid acidification of the apoplast via the activation of PM-resident P-type proton pumps (Caesar et al., 2011; Grosseholz et al., 2022; Witthöft et al., 2011)

RLP44 is proposed to be a cell wall integrity sensor that controls cell wall homeostasis by interplay with BRI1 and BAK1 (Wolf et al., 2012; Wolf et al., 2014). In fact, we were recently able to demonstrate the existence of a ternary RLP44/BRI1/BAK1 complex in the PM of living plant cells using a spectral Förster resonance energy transfer (FRET) and FRET-Fluorescence-lifetime imaging microscopy (FLIM) approach (Glöckner et al., 2022). Additionally, RLP44 has been linked to phytosulfokine signaling. The corresponding receptor Phytosulfokine Receptor 1 (PSKR1) is also proposed to form a complex with RLP44 and BAK1 (Gomez et al., 2021; Holzwart et al., 2018).

### OneFlowTraX analysis of BRI1 and RLP44 dynamics and nano-structured organization in the Nicotiana benthamiana transient expression system and transgenic Arabidopsis seedlings

For initial assessment, mEos3.2, PA-GFP and PATagRFP-tagged BRI1 fusion proteins were transiently expressed in *Nicotiana benthamiana* (*N. benthamiana*) epidermal leaf cells. Such transient expression setups provide a fast and convenient way to test the functionality of protein fusions with photoswitchable/photoconvertible fluorophores (Gronnier et al, 2017; Perraki et al, 2018). For the transgenic approach in *A. thaliana,* the mEos3.2, PA-GFP and PATagRFP-tagged fusions of RLP44 were chosen.

The density of the single fusion proteins after photoconversion or –activation was optimal (Figure 5A) for a comprehensive evaluation of sptPALM data (Bayle et al, 2021).

**Figure 5.**
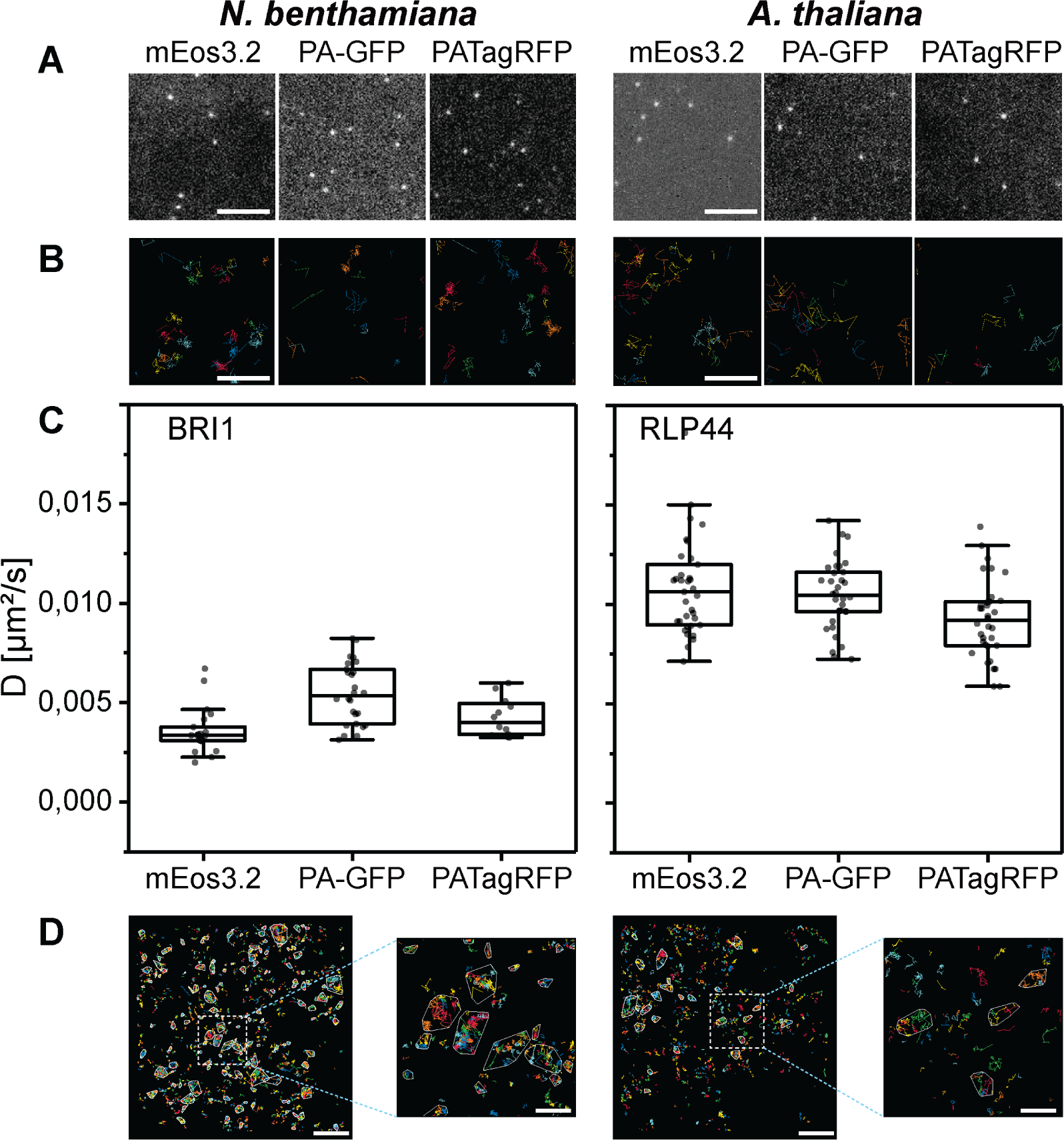
BRI1 and RLP44 FP fusions accumulate in the PM of transiently transformed *N. benthamiana* epidermal leaf cells or transgenic *A. thaliana* epidermal hypocotyl cells, respectively, and their nanoscale organization can be evaluated by OneFlowTraX. (A) Representative images from sptPALM movies, showing, from left to right, single mEos3.2, PA-GFP and PATagRFP of BRI1 in transiently transformed *N. benthamiana* leaf epidermal cells and of RLP44 in light-grown transgenic *A. thaliana* hypocotyl epidermal cells. The BRI1 and RLP44 fusion proteins were expressed via their native promoters (pBRI1, pRLP44). Scale bar = 1 µM. **(B)** Representative magnifications of areas containing tracks generated by OneFlowTraX (random coloring per track) after localization of the BRI1 and RLP44 fusion proteins, with panels ordered as above. Scale bar = 100 nm. **(C)** Diffusion coefficients for the indicated BRI1 and RLP44 fusion proteins in the PM obtained with OneFlowTraX. For statistical evaluation, the data were checked for normal distribution and unequal variances and then analyzed applying the Kruskal-Wallis test followed by the Steel-Dwass post-hoc test, with n ≥ 12 for each fusion protein derived from at least five plants, measured on at least three different days. Whiskers show the data range excluding outliers, while the boxes represent the 25-75 percentile. For detailed statistics see Supplemental Table 3. **(D)** The track data was used for cluster analysis in OneFlowTraX, exemplified for BRI1-mEos3.2 and RLP44-mEos3.2 clusters that were assigned using the NASTIC algorithm. The images represent one cell with random coloring per track and cluster boundaries highlighted in white. Scale bar = 2 µm (500 nm for the magnified view).

Using OneFlowTraX (for detailed parameters see Supplemental Table 4), the data were processed, and single-molecule trajectories were generated (Figure 5B). After MSD analysis, the diffusion coefficients were calculated (Figure 5C).

Although the three fluorophores are sptPALM-optimized versions of different precursors from various marine organisms, the BRI1 fusion proteins showed comparable diffusion coefficients in *N. benthamiana* epidermal leaf cells (Figure 5C, left panel). The same was observed for the RLP44 fusion proteins in transgenic *Arabidopsis* epidermal hypocotyl cells (Figure 5C, right panel). This demonstrates that all three fluorophores can be reliably used for sptPALM studies in different plant cell systems. The best signal-to-noise ratio was obtained with mEos3.2, due to its excellent photophysical properties (Supplemental Table 1). On the other hand, PA-GFP and PATagRFP can be combined - due to their non-overlapping spectra - for dual-color sptPALM experiments.

Moreover, as shown in Figure 5D, OneFlowTraX allows the use of sptPALM single-molecule track data to show that the BRI1-mEos3.2 and RLP44-mEos3.2 fusion proteins are partially organized in clusters.

### OneFlowTraX application examples to detect changes in protein mobility

To validate the usefulness of OneFlowTraX, we first analyzed the dynamics of LTi6a-mEos2 and PIP2;1-mEos2 (Hosy et al, 2015) in the PM of epidermal root tip and epidermal hypocotyl cells of transgenic *A. thaliana* seedlings. In accordance with their previous results, we observed that LTi6a exhibited a significantly higher mobility than PIP2;1 in the PM of root epidermal cells (Figure 6A, left). We found that this difference was even more pronounced in the PM of epidermal hypocotyl cells (Figure 6A, right). Moreover, the mobility of LTi6a was significantly higher in the PM of epidermal hypocotyl cells than in epidermal cells of the root. In contrast, we observed no difference for PIP2;1 (Figure 6A).

**Figure 6.**
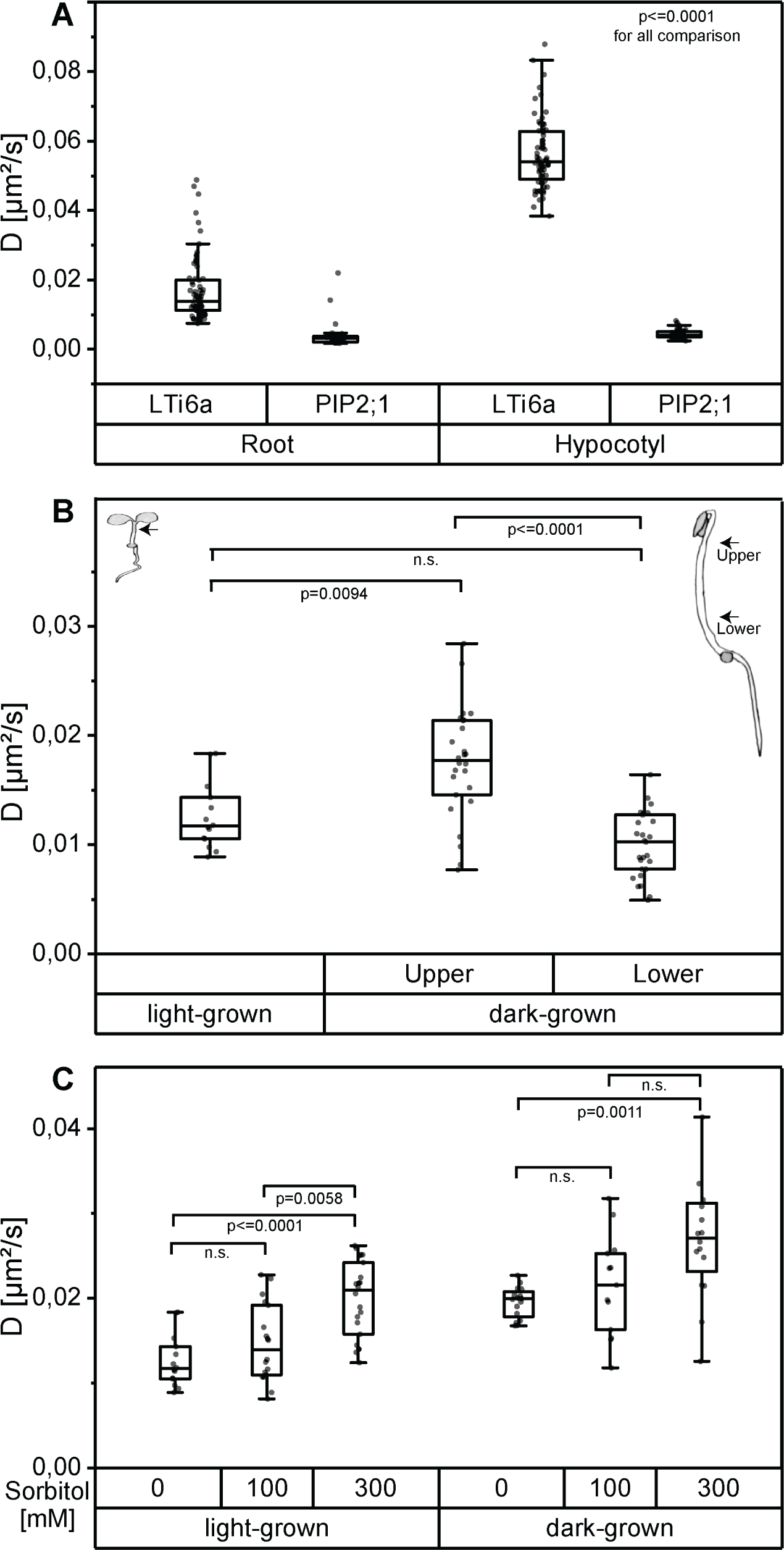
OneFlowTraX analysis of sptPALM data reveals organ- and position-specific as well as environmentally caused differences in the dynamics of the indicated FP fusion proteins in epidermal cells of *A. thaliana*. (**A**) Diffusion coefficients of *pPIP2;1* expressed LTi6a-mEos2 and PIP2;1-mEos2 in epidermal root and hypocotyl cells of light-grown *A. thaliana* seedlings. For statistical evaluation, the data were checked for normal distribution and unequal variances and then analyzed applying the Kruskal-Wallis test followed by the Steel-Dwass post-hoc test, with n ≥ 37 for each fusion protein derived from at least six plants, measured on at least four different days. Whiskers show the data range excluding outliers, while the boxes represent the 25-75 percentile. For detailed statistics see Supplemental Table 3. **(B)** Diffusion coefficients of *pRLP44* expressed RLP44-mEos3.2 in epidermal hypocotyl cells of light-grown and dark-grown *A. thaliana* seedlings. The locations of the recorded measurements are indicated by arrows in the corresponding seedling sketches. For dark-grown seedlings, the center of the distance from the apical hook to the seed was used as the reference point. Based on this, the centers to the apical hook and to the seed were used as selected regions, named ‘upper’ and ‘lower’, respectively. Statistical evaluation was performed as described in **(A)** with n ≥ 14, from at least three plants. For detailed statistics see Supplemental Table 3. **(C)** Diffusion coefficients of *pRLP44* expressed RLP44-mEos3.2 recorded from epidermal hypocotyl cells of light-grown or dark-grown *A. thaliana* seedlings after 20 min treatment with sorbitol solution of 100 mM or 300 mM or after mock-treatment (0 mM). For the recording of the data in the dark-grown seedlings the upper part of the hypocotyl cells was chosen (see **(B)**). Statistical evaluation was performed as described in (A), with n ≥ 13, from three different plants. For detailed statistics see Supplemental Table 3.

We then conducted OneFlowTraX analysis of the RLP44-mEos3.2 dynamics in transgenic *A. thaliana* seedlings grown either in white light or darkness in order to determine whether the developmental state (photomorphogenic versus skotomorphogenic) influences the fusion protein’s dynamics. Firstly, as shown in Figure 6B, RLP44-mEos3.2 moved significantly faster in the PM of cells of the upper part of dark-grown hypocotyls compared to the lower part. The diffusion coefficient of RLP44-mEos3.2 in the PM of hypocotyl cells of light-grown seedlings, where a differentiation of different cell types along the axis is hardly possible, was similar to the diffusion coefficients gained from the lower part of the etiolated hypocotyl. These in planta generated results show that positional and developmental differences in the RLP44-mEos3.2 dynamics can be detected by sptPALM-based OneFlowTraX analysis.

The effect of other environmental factors on protein mobility can also be evaluated by OneFlowTraX. Here we chose the hyper-osmotic stress response of plant cells after addition of sorbitol, a non-toxic, widely used, non-metabolic osmolyte already applied for sptPALM (Hosy et al, 2015). Sorbitol induces a cell volume loss (plasmolysis) when used at a concentration of 300 mM. Under milder hyper-osmotic conditions (100 mM), the cells experience a reduction in turgor without major changes in their volume (Martiniere et al., 2019). Thus, the increasing concentration of sorbitol from 100 to 300 mM induces a progressive separation of the PM from the cell wall which enhanced the dynamics of almost all intrinsic PM proteins (Hosy et al, 2015). Based on this knowledge, treatments with 100 mM and 300 mM sorbitol or mock treatment (all applied for 20 min) were performed with light-grown and dark-grown seedlings expressing RLP44-mEos3.2. Then, the sptPALM data were measured in epidermal hypocotyl cells of light-grown seedlings and epidermal cells of the upper hypocotyl part of dark-grown seedlings and analyzed using OneFlowTraX. Whereas the concentration of 100 mM sorbitol did not significantly affect the diffusion coefficient of RLP44-mEos3.2 compared to the mock treatment (0 mM), 300 mM sorbitol significantly increased the protein mobility in all measured cells (Figure 6C).

These examples represent common “blueprints” for the analyses of sptPALM data in plant research and demonstrate the applicability of OneFlowTraX, as it was used to perform all evaluation steps, including localization, tracking, protein mobility and cluster analysis.

## DISCUSSION

### OneFlowTraX is an all-in-one software package for SMLM data processing and analysis

With OneFlowTraX, we provide a unique, user-friendly open-source software package that guides users through the steps of localization, tracking, analysis of mobility and nanoscale organization of single molecules based on SMLM data. OneFlowTraX facilitates the previously very time-consuming and cumbersome post-acquisition processing of SMLM data. Although OneFlowTraX is tailored for use by cell biologists, who want to perform sptPALM analyses in the PM of plant cells, it also offers the possibility of post-acquisition data processing for other non-PM bound processes in any prokaryotic and eukaryotic cells. For instance, the software could be used to analyze the spatial organization of chromosomes into topologically associating domains (Wang et al., 2016).

In plants, due to the cell wall, only regions close to the microscope slide can be analyzed such as the PM and PM-associated molecules. OneFlowTraX provides a default set of analysis parameters that are well suited for plant PM molecules located in a 2D environment. However, as described in the provided manual, all settings can be adjusted by the user to their specific experimental requirements for the SMLM analysis of any process of interest independent of the organism.

The primary objective of OneFlowTraX is to integrate established algorithms from the community into a pipeline and to optimize data flow. We have successfully combined approaches from specialized software that, for example, either focuses on single-molecule localization or tracking. Our integrated solution also enables the utilization of track data for cluster analyses, which is particularly important in single-molecule tracking experiments. OneFlowTraX includes comprehensive batch analysis, with detailed parameter and data output for all analysis paths, a feature that was previously missing from many analysis tools. This not only saves the user time, but also maintains the consistency of the single-molecule data with the calculated results, making it easier to apply the FAIR principle. The detailed but clearly structured parameter organization also provides a method to evaluate the reliability of the analyses: running multiple batch analyses with different parameters will show the impact of individual parameters in the overall analysis process on the results.

OneFlowTraX was programmed in MATLAB, a software language that is easy to work with and widely used in academia. The code has been designed in a clear and concise manner, with comments provided to aid in the insertion of changes or extensions. This could also make OneFlowTraX of interest to scientists working with animal or human cells.

### PATagRFP, PA-GFP and mEos3.2 are suitable FPs for sptPALM analyses in living plant cells

Especially plant scientists rely on genetically encoded photoactivatable or photoconvertible FPs fused to the protein of interest as the cell wall is impenetrable for dyes applied from the outside. With mEos3.2, PA-GFP and PATagRFP, three different FPs are now available that do not tend to aggregate, deliver satisfactory signal-to-noise ratios during imaging and are therefore applicable for sptPALM applications both in a transient transformation system (*N. benthamiana*) and in stably transformed *A. thaliana* plants. We expect that these fluorophores will also work in other plant species. Of particular interest is the combination of PA-GFP and PATagRFP, which will enable dual-color sptPALM in the future for simultaneous recording and spatiotemporal analysis of two different fusion proteins. Because of its superior photophysical properties, we recommend the use of mEos3.2 for single FP fusion protein analyses. Especially when analyzing the mobility of PM proteins in living plant cells, the sptPALM technology shows its superiority in terms of spatiotemporal resolution compared with other methods such as FRAP and FCS (Bayle et al, 2021).

### OneFlowTraX allows the rapid in vivo analysis of PM proteins in transiently and stably transformed plant cells

We used two functionally well-studied PM proteins, namely BRI1 and RLP44, for our proof-of-principle experiments regarding the usability of the three FPs and the applicability of OneFlowTraX. Both proteins were evaluated in either transiently transformed *N. benthamiana* epidermal leaf cells or the epidermis of the hypocotyl of stable transgenic *A. thaliana* seedlings. Importantly, the accumulation of the BRI1 and RLP44 fusion proteins was driven by the respective endogenous promoters. Our experience has been that strong over-accumulation of the fusion proteins driven by constitutively active promoters leads to overcrowded sptPALM movies in both plant systems. Analysis of these movies results in misconnected tracks that affect the entire OneFlowTraX analysis causing inadequate results in subsequent steps. Therefore, we recommend using weaker promoters whenever possible.

While OneFlowTraX further demonstrated its utility through the replication of LTi6a-mEos2 and PIP2;1-mEos2 sptPALM-derived findings in the roots of light-grown *A. thaliana* seedlings (Hosy et al, 2015), we were additionally able to observe that the PM dynamics of both mEos2 fusions differ between epidermal cells of the root and hypocotyl. This suggests that the organ context of a tissue has an influence on PM protein dynamics in plant cells.

Moreover, during our OneFlowTraX analysis of the RLP44-mEos3.2 dynamics in epidermal cells of light-grown *Arabidopsis* hypocotyls, we observed high data variability. This led to the hypothesis that cell-specific effects depending on the cell’s position in the organ, i.e., on its physiological state, interfere with the membrane dynamics of the fusion protein to be investigated. This hypothesis was substantiated by the technically easier access to the potential positional effects in the dynamics of RLP44-mEos3.2 in epidermal cells along the hypocotyl axis of dark-grown seedlings: The diffusion coefficient of RLP44-mEos3.2 is significantly higher in the epidermal cells of the upper part than in those of the lower part of the dark-grown hypocotyl. Furthermore, the diffusion coefficient in the lower part is similar to that found in the epidermal cells of light-grown hypocotyls. Thus, OneFlowTraX can easily capture differences in dynamics, allowing SMLM in different cells and tissues that may vary in their membrane properties.

As previously shown by Hosy et al, 2015 and Martiniere et al, 2019, the treatment of *A. thaliana* root cells with increasing concentration of the osmolyte sorbitol increases the diffusion coefficient of membrane proteins such as the aquaporin PIP2;1. Using OneFlowTraX, we could substantiate these findings for the RLP44-mEos3.2 fusion and demonstrated potential effects of the differentiation states of the analyzed tissue. This shows that reproducible and robust SMLM data analysis is provided by OneFlowTraX.

In addition to the analysis of protein dynamics, OneFlowTraX also offers different algorithms (Voronoi tessellation, DBSCAN, NASTIC) for the analysis of the nanoscale organization of a given membrane protein. For sptPALM data, we recommend the NASTIC algorithm as it is specifically designed to work with track data. Because of the sensitivity to parameter changes, care must be taken when regarding the results of the nanoscale evaluation output by all of these algorithms as absolute values. However, relative comparisons of the nanoscale organization for a given membrane protein are possible if there are no changes in the parameter settings between the experiments.

In summary, SMLM data acquisition such as that from sptPALM becomes more easily accessible and faster analyzable with OneFlowTraX. The fluorophores mEos3.2, PA-GFP and PATagRFP have proven to be suitable for SMLM in planta and allow analysis of membrane proteins in transient expression systems as well as in stable transformed plants. The respective FP fusions are suitable for the analysis of protein dynamics in epidermal cells of different organs and at different developmental or physiological stages as well as in response to environmental factors. Therefore, OneFlowTraX will greatly facilitate the comprehensive investigation of the dynamics and nanoscale organization of single molecules in the future.

## MATERIALS AND METHODS

### Plasmid construction

All expression clones were constructed using GoldenGate assembly with BB10 as the vector (Binder et al., 2014). Promoter sequences were obtained with the help of the Integrated Genome Browser (Freese et al., 2016). Level I modules were generated by PCR amplification of the desired sequences and then blunt-end cloned into pJET1.2 (Thermo Fisher Scientific). Fluorophores were designed as C-terminal fusions (D-E module) using either a glycine/serine or a glycine/alanine-rich linker. The coding sequences of BRI1 and RLP44 were constructed as B-D modules, eliminating the need for a B-C dummy module. A full list of used constructs can be found in Supplemental Table 5. The correctness of Level I constructs was confirmed by Sanger sequencing. Cut-ligations for the Level II generation were performed with 40 cycles, without using bovine serum albumin as described by Binder et al, 2014. Reactions were transformed into TOP10 cells (Thermo Fisher Scientific), and colony correctness was verified via restriction enzyme analysis and partial Sanger sequencing.

### Plant material and growth conditions

The transgenic *A. thaliana* lines generated for this study were all in the Columbia (Col-0) background. The respective stable lines were created using the Floral dip method according to Zhang et al., 2006. For the reproduction of LTi6a and PIP2;1 results, seeds of the corresponding lines were provided by Dr. Doan-Trung Luu. Transgenic seeds were propagated either based on the presence of the pFAST marker by binocular visual inspection or by selection of survivors on ½ Murashige and Skoog (MS) plates containing 1 % (w/v) sucrose and 0.8 % (w/v) phytoagar supplemented with 25 µM hygromycin. For sptPALM measurements, seeds were sterilized with a solution of 70 % ethanol (v/v) and 0.05 % Triton X-100 for 30 minutes followed by a 10-minute treatment with absolute ethanol. Seeds were sown on ½ MS plates (+1 % sucrose and 0.8 % phytoagar) and stratified at 4 °C for at least 24 hours. For measurements of dark-grown seedlings, seeds were exposed to light from the growth chamber for two hours before being wrapped in aluminum foil until the day of measurement. Light-grown seedlings were cultivated in growth chambers at 20 °C under long-day conditions (16 hours light / 8 hours dark). The duration of growth is indicated in the respective figures. The *N. benthamiana* plants used in this study were cultivated under controlled greenhouse conditions. Proteins were transiently expressed using the AGL1 *Agrobacterium tumefaciens* strain (Lifeasible), as previously described (Hecker et al., 2015; Ladwig et al., 2015), without the washing step with sterile water. The plants were infiltrated with the respective construct at an OD600 of 0.1, in a ratio of 1:1 with the silencing inhibitor p19. After watering, the plants were kept in ambient conditions and were imaged three days after infiltration.

### Sample preparation and movie acquisition

All sptPALM measurements with transiently transformed *N. benthamiana* were performed three days after infiltration. A small leaf area was cut out, excluding veins, and placed between two coverslips (Epredia 24×50 mm #1 or equivalent) with a drop of water. This “coverslip sandwich” was then placed on the specimen stage, lightly weighted down by a brass ring to help flatten the uneven cell layers, especially in *N. benthamiana*. Seven-day-old *A. thaliana* seedlings were used to acquire data from stable *Arabidopsis* lines. Depending on the experiment, either light-grown or dark-grown plants were used. For sorbitol (obtained by Roth) treatments, seedlings were incubated with the appropriate concentration in 12-well plates for five minutes before being transferred to the coverslip and imaged in the respective incubation solution as mounting medium for up to 20 minutes. Similar to the handling of *N. benthamiana* leaf discs, the seedlings were placed between coverslips and brass rings.

The custom-built microscope platform for sptPALM acquisition is described in detail in Supplementary Materials and Methods. Briefly, lasers of different wavelengths and their intensities are controlled by an acousto-optic transmission filter. A laterally translatable lens in the excitation beam path allows to adjust the VAEM illumination of the sample utilizing a high NA objective. The emitted light from the sample is separated from the excitation light by a multi-band beam splitter and is detected by an sCMOS camera. Depending on the fluorophore fusion, the following filters were inserted into the emission beam path: (i) mEos3.2: 568 LP Edge Basic Longpass Filter, 584/40 ET Bandpass; (ii) PA-GFP: 488 LP Edge Basic Longpass Filter, 525/50 BrightLine HC; (iii) PATagRFP: 568 LP Edge Basic Longpass Filter, 600/52 BrightLine HC (all AHF analysentechnik AG). The excitation power arriving at the sample was measured (PM100D with S120C, Thorlabs) in epifluorescence mode after the objective to keep it constant for the respective experiment sets. If necessary, photoconversion or photoactivation was performed using 405 nm excitation at varying low intensities (for detailed acquisition parameters see Supplemental Table 4). The magnification of the optical system was adjusted so that the length of one camera pixel corresponds to 100 nm in the sample plane. Viable regions of interest were screened in a larger area of 51.2 x 51.2 µm by adjusting the focal plane and the VAEM angle with a frame rate of 10 Hz, while recording was performed with 12.8 x 12.8 µm and frame rates between 20 and 50 Hz, recording between 2 500 and 5 000 frames per movie (see Supplemental Table 4). For each measurement day, noise files (a series of dark images) were recorded with the corresponding frame rates.

#### Raw data processing and analysis with OneFlowTraX

Subsets of each experimental data set were loaded into OneFlowTraX to inspect the quality of the data and to find appropriate parameters for each analysis step as described above. On the Batch Analysis tab, all sptPALM raw data files that share the same parameter set were processed together (see Supplemental Table 4 for a detailed list of applied analysis parameters). Samples that showed significantly low numbers of localizations or tracks compared to others in the same batch were discarded.

## SUPPLEMENTAL INFORMATION

All Supplemental information is provided with this document (see below).

## AUTHOR CONTRIBUTIONS

Conceptualization, A.J.M., B.K., K.H., and S.z.O.-K.; Software, S.z.O.-K.; Formal Analysis, L.R. and S.z.O.-K.; Data Curation, L.R., Ll.R., and S.z.O.-K.; Methodology, S.z.O.-K.; Investigation, L.R., Ll.R., and S.z.O.-K.; Writing – Original Draft, K.H., L.R., and S.z.O.-K.; Writing – Review & Editing, A.E., A.J.M., B.K., K.H., J.G., L.R., Ll.R., N.G.B., and S.z.O.-K.; Visualization, L.R. and S.z.O.-K.; Funding Acquisition, A.J.M., B.K., and K.H.; Resources, A.E., A.J.M., and N.G.B.; Supervision, A.J.M., J.G., B.K., and K.H.

## DATA AND SOFTWARE AVAILABILITY

The OneFlowTraX software (including a user guide), is freely available to non-commercial users at GitHub (https://github.com/svenzok/OneFlowTraX).

## ACKNOWLEDGMENTS

We would like to thank Dr. Andrea Gust for providing us with the pFAST construct. We would also like to thank Dr. Doan-Trung Luu for providing seeds of the LTi6a-mEos2 and PIP2;1-mEos2 transgenic *A. thaliana* lines. No conflict of interest is declared.

## SUPPLEMENTAL INFORMATION

**This supplemental information file contains:**

**Supplementary Tables 1-5**

**Supplementary Figure 1**

**Supplementary Materials and Methods**

## SUPPLEMENTARY TABLES

**Supplemental Table 1.** Photophysical characteristics of the fluorophores mEos2, mEos3.2, PA-GFP and PATagRFP. For the mEos fluorophores, both the native green (G) and the photoconverted red (R) form are listed, while for PA-GFP and PATagRFP, only the activated forms are presented. The molar attenuation coefficient (ε) refers to the excitation maximum, brightness is the product QY·ε (for comparison: EGFP features a brightness of 33.6 mM^-1^ cm^-1^). Mean label density was compared in HeLa cells. Note that according to the referenced publication, the high label density of mEos3.2 (compared with mEos2) is likely due to its better maturation and folding properties.

|  | mEos2<br>(G) | mEos2<br>(R) | mEos3.2<br>(G) | mEos3.2<br>(R) | PA-GFP | PAtagRFP |
| --- | --- | --- | --- | --- | --- | --- |
| Excitation maximum (nm) | 506 | 573 | 507 | 572 | 504 | 562 |
| Emission maximum (nm) | 519 | 584 | 516 | 580 | 517 | 595 |
| Quantum yield | 0.84 | 0.47 | 0.84 | 0.55 | 0.79 | 0.38 |
| $\epsilon$ ($\text{M}^{-1} \text{ cm}^{-1}$) | 56 000 | 41 300 | 63 400 | 32 200 | 17 400 | 66 000 |
| Brightness ($\text{mM}^{-1} \text{ cm}^{-1}$) | 47 | 19 | 53 | 18 | 14 | 25 |
| pH stability ($\text{pK}_a$) | 5.6 | 6.0 | 5.4 | 5.8 | n.d. | 5.3 |
| Photostability $t_{1/2}$ (s) | 14.0 | 48.0 | 12.6 | 48.0 | n.d. | 180 |
| Oligomerisation $K_d$ ($\mu\text{M}$) | 20 | 20 | - | - | - | - |
| Mean label density ($\mu\text{m}^{-2}$) | n.d. | 2 649 | n.d. | 9 865 | n.d. | n.d. |
| Reference for measured values | (Zhang et al., 2012) | (Zhang et al., 2012) | (Zhang et al., 2012) | (Zhang et al., 2012) | (Patterson and Lippincott-Schwartz, 2002) | (Subach et al., 2010) |

**Supplemental Table 2.**
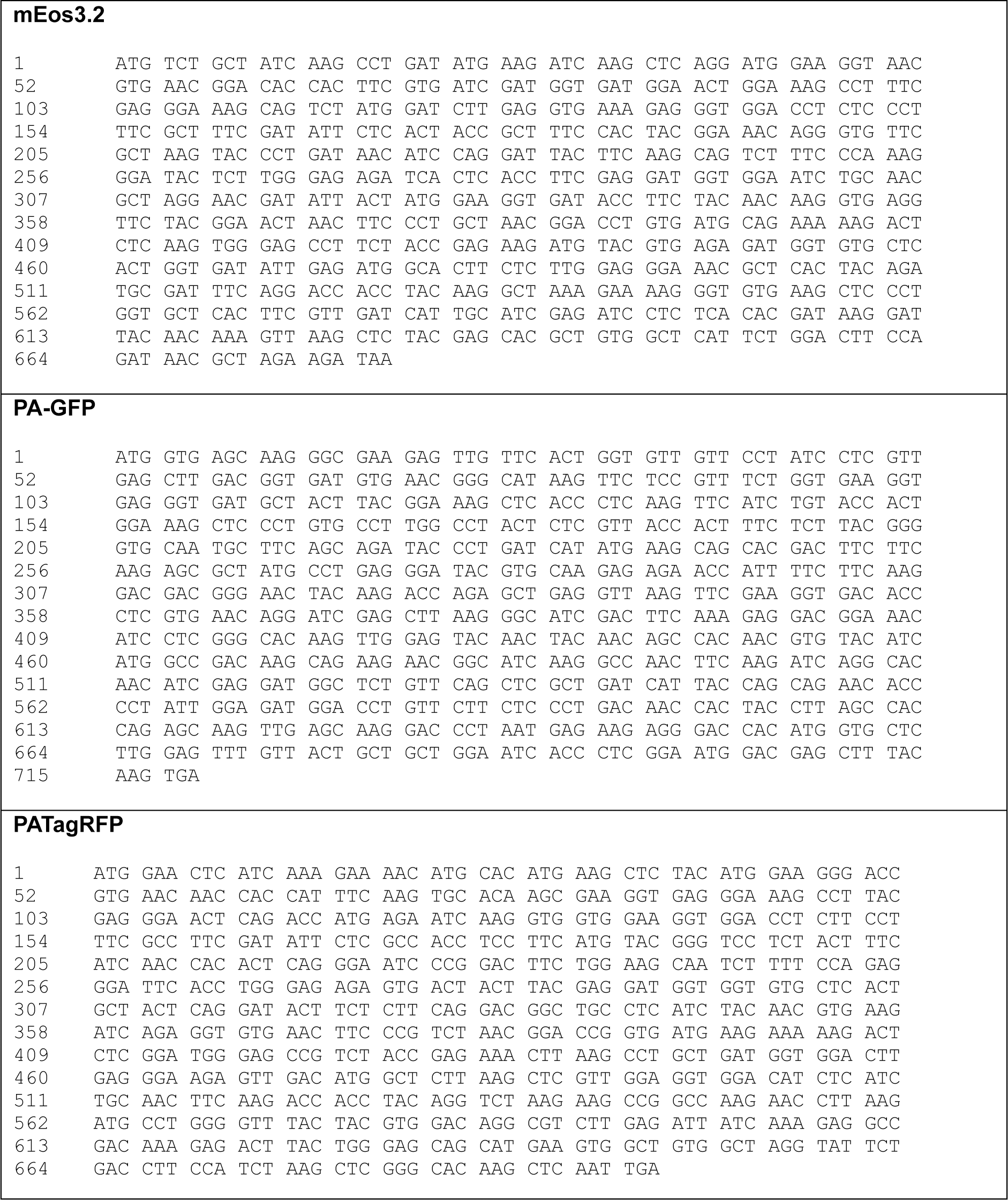
Nucleotide sequences of the codon-optimized fluorophores mEos3.2, PA-GFP and PATagRFP. Shown are the codon optimized nucleotide sequences of the fluorophores mEos3.2, PA-GFP and PATagRFP and the respective position. Genes were synthesized by Invitrogen’s GeneArt services (Thermo Fisher Scientific)

**Supplemental Table 3.**
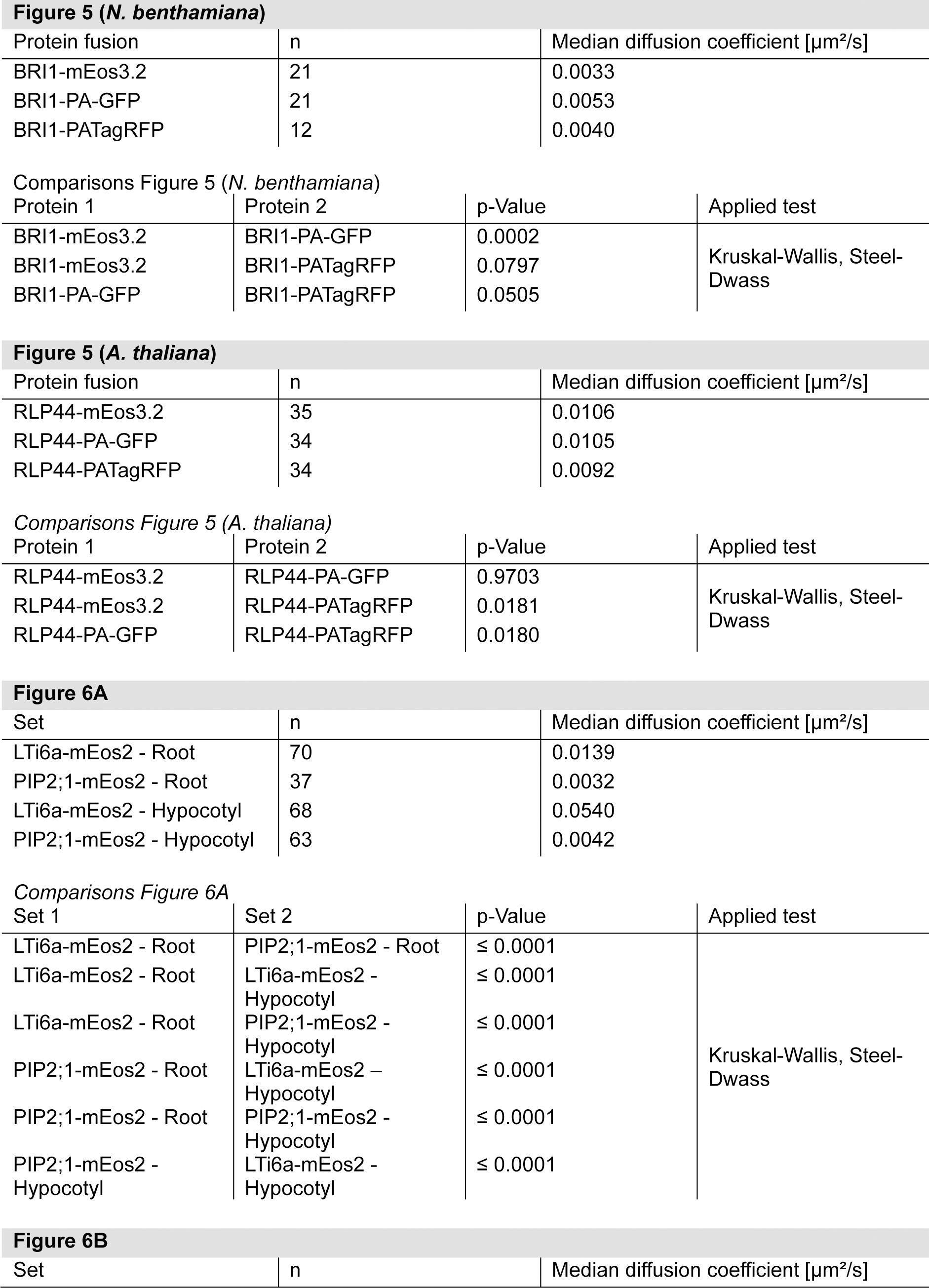
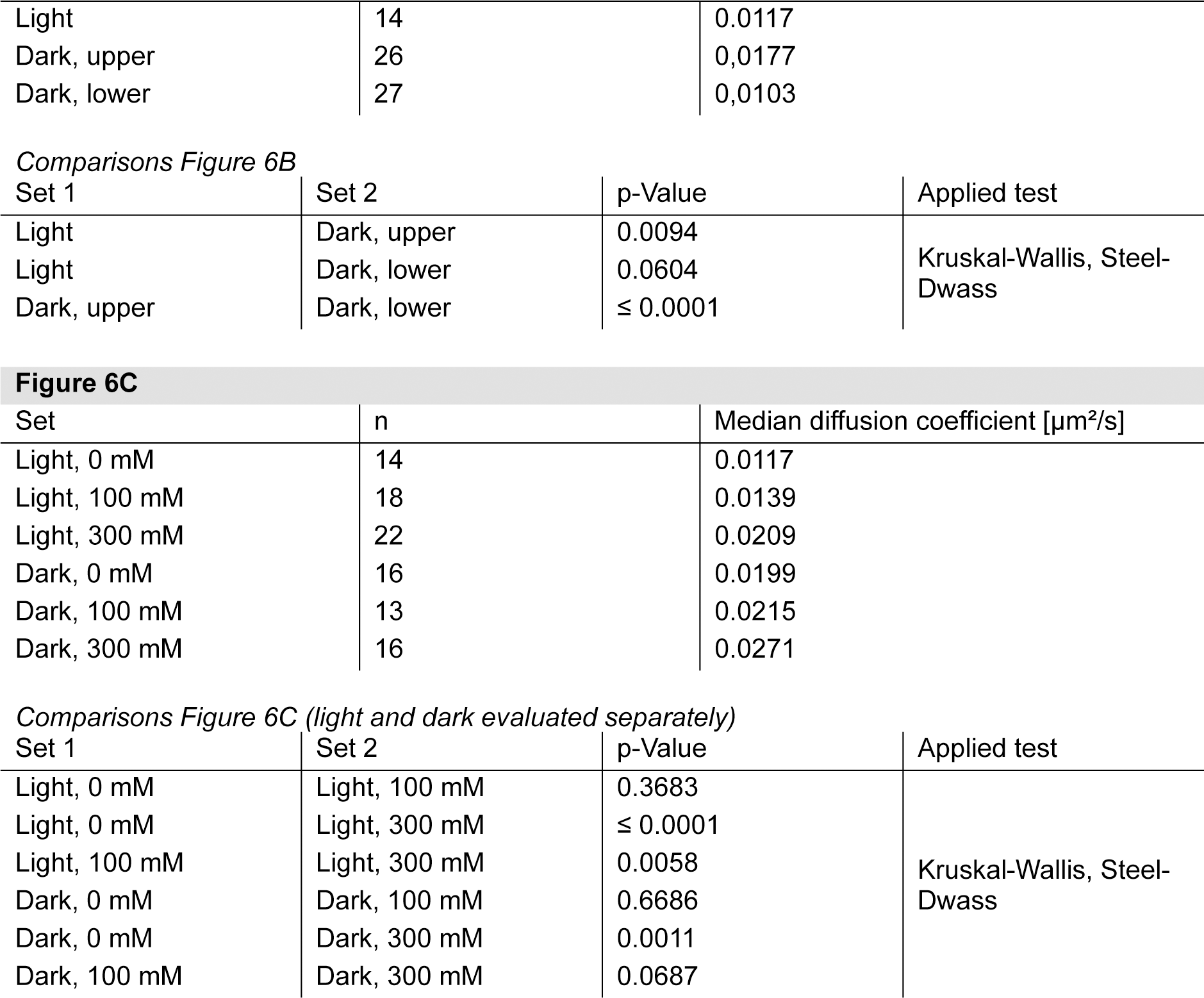
Statistical information about applied tests for the respective figures. All protein fusions or treatment conditions are listed with the number of replicates (n) and the median diffusion coefficient (in µm²/s). In addition, all compared combinations and the corresponding p-value based on the applied test, are shown.

**Supplemental Table 4.** List of sptPALM acquisition and analysis parameters.

|  | *N. benthamiana* constructs with |  |  | *A. thaliana* constructs with |  |  |
| --- | --- | --- | --- | --- | --- | --- |
|  | mEos3.2 | PA-GFP | PATagRFP | mEos3.2 | PA-GFP | PATagRFP |
| acquisition |  |  |  |  |  |  |
| Power 405 nm (μW) | <15 | <15 | <15 | <20 | <20 | <40 |
| Power 488 nm (μW) |  | 600 |  |  | 900 |  |
| Power 561 nm (μW) | 1800 |  | 1300 | 1800 |  | 2000 |
| frame rate (Hz) | 20 | 20 | 20 | 50 | 20 | 20 |
| number of frames | 2500-5000 | 2500-5000 | 2500-5000 | 5000 | 2500 | 2500 |
| localization |  |  |  |  |  |  |
| conversion | 0.48 | 0.48 | 0.48 | 0.48 | 0.48 | 0.48 |
| offset | 100 | 100 | 100 | 100 | 100 | 100 |
| pixel size | 100 | 100 | 100 | 100 | 100 | 100 |
| filter size | 1.2 | 1.2 | 1.2 | 1.2 | 1.3 | 1.3 |
| cut-off | 3 | 3 | 3 | 3 | 3.5 | 4 |
| PSF ROI size | 7 | 7 | 7 | 7 | 7 | 7 |
| tracking |  |  |  |  |  |  |
| max. linking distance | 200 | 200 | 200 | 200 | 200 | 200 |
| max. gap frames | 4 | 4 | 4 | 4 | 4 | 4 |
| mobility analysis |  |  |  |  |  |  |
| min. number of datapoints | 8 | 8 | 8 | 8 | 8 | 8 |
| number of points for linear fit | 4 | 4 | 4 | 4 | 4 | 4 |
| adj. R<sup>2</sup> fit threshold value | 0.4 | 0.4 | 0.4 | 0.4 | 0.4 | 0.4 |

**Supplemental Table 5.**
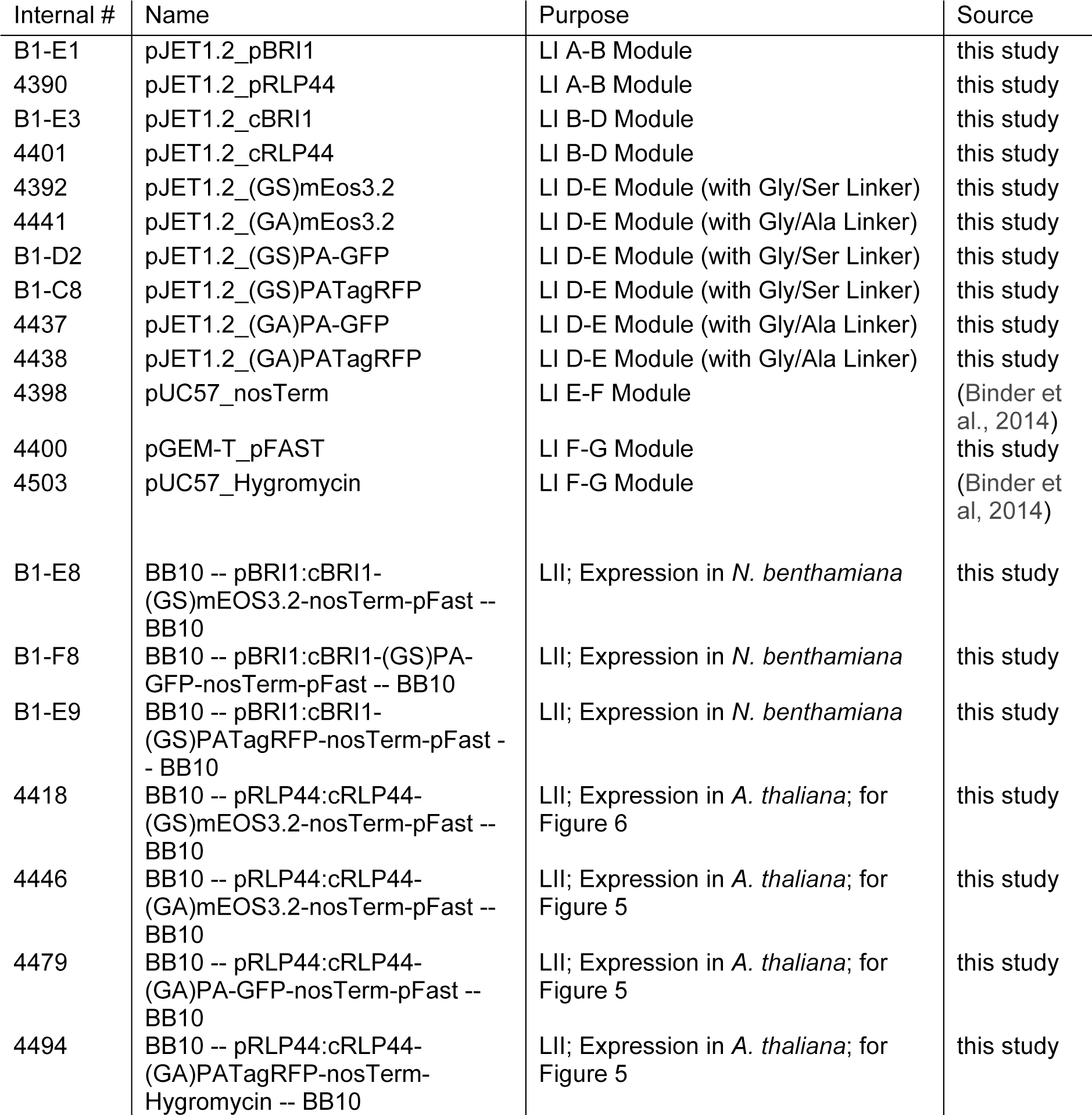
List of constructs. All used constructs are listed with our internal numbering, their name, their purpose and source.

| Internal # | Name | Purpose | Source |
| --- | --- | --- | --- |
| B1-E1 | pJET1.2_pBRI1 | LI A-B Module | this study |
| 4390 | pJET1.2_pRLP44 | LI A-B Module | this study |
| B1-E3 | pJET1.2_cBRI1 | LI B-D Module | this study |
| 4401 | pJET1.2_cRLP44 | LI B-D Module | this study |
| 4392 | pJET1.2_(GS)mEos3.2 | LI D-E Module (with Gly/Ser Linker) | this study |
| 4441 | pJET1.2_(GA)mEos3.2 | LI D-E Module (with Gly/Ala Linker) | this study |
| B1-D2 | pJET1.2_(GS)PA-GFP | LI D-E Module (with Gly/Ser Linker) | this study |
| B1-C8 | pJET1.2_(GS)PATagRFP | LI D-E Module (with Gly/Ser Linker) | this study |
| 4437 | pJET1.2_(GA)PA-GFP | LI D-E Module (with Gly/Ala Linker) | this study |
| 4438 | pJET1.2_(GA)PATagRFP | LI D-E Module (with Gly/Ala Linker) | this study |
| 4398 | pUC57_nosTerm | LI E-F Module | (Binder et al., 2014) |
| 4400 | pGEM-T_pFAST | LI F-G Module | this study |
| 4503 | pUC57_Hygromycin | LI F-G Module | (Binder et al., 2014) |
| B1-E8 | BB10 -- pBRI1:cBRI1-(GS)mEOS3.2-nosTerm-pFast -- BB10 | LII; Expression in *N. benthamiana* | this study |
| B1-F8 | BB10 -- pBRI1:cBRI1-(GS)PA-GFP-nosTerm-pFast -- BB10 | LII; Expression in *N. benthamiana* | this study |
| B1-E9 | BB10 -- pBRI1:cBRI1-(GS)PATagRFP-nosTerm-pFast -- BB10 | LII; Expression in *N. benthamiana* | this study |
| 4418 | BB10 -- pRLP44:cRLP44-(GS)mEOS3.2-nosTerm-pFast -- BB10 | LII; Expression in *A. thaliana*; for Figure 6 | this study |
| 4446 | BB10 -- pRLP44:cRLP44-(GA)mEOS3.2-nosTerm-pFast -- BB10 | LII; Expression in *A. thaliana*; for Figure 5 | this study |
| 4479 | BB10 -- pRLP44:cRLP44-(GA)PA-GFP-nosTerm-pFast -- BB10 | LII; Expression in *A. thaliana*; for Figure 5 | this study |
| 4494 | BB10 -- pRLP44:cRLP44-(GA)PATagRFP-nosTerm-Hygromycin -- BB10 | LII; Expression in *A. thaliana*; for Figure 5 | this study |

## SUPPLEMENTARY FIGURES

**Supplemental Figure 1.**
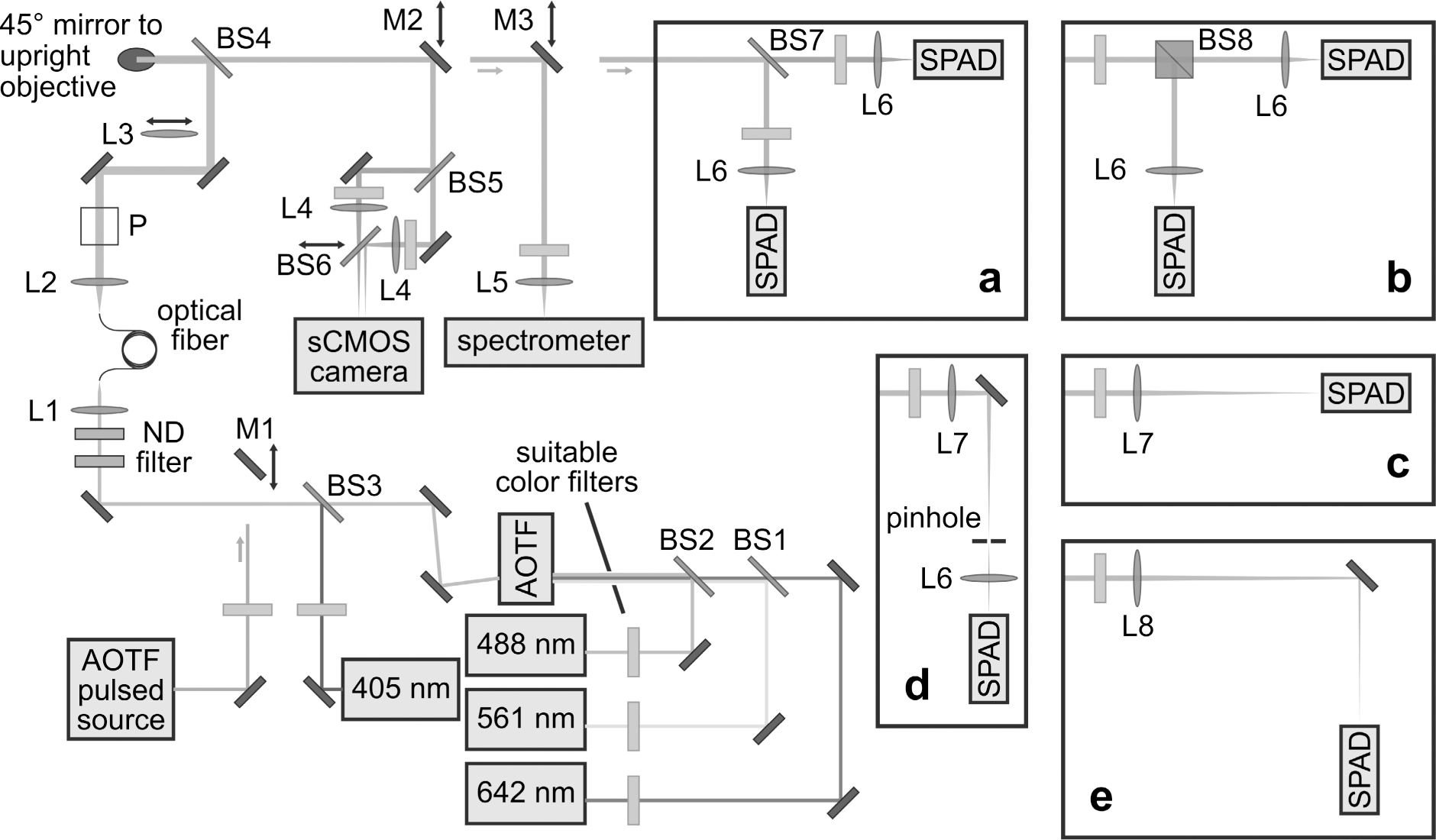
Compact schematic overview of the research microscope. For a detailed description and references to numbered items like lenses (L), beam splitters (BS) and mirrors (M) see text in Supplementary Materials and Methods, also for the different confocal configurations depicted in insets (a-e). Abbreviations: Acousto-optical tuneable filter (AOTF), neutral density filter (ND filter), sCMOS (scientific complementary metal-oxide-semiconductor) camera, single-photon avalanche diode (SPAD). The box P in the beam path denotes the position for additional elements, such as polarization optics. Adapted from (zur Oven-Krockhaus, 2021).

## SUPPLEMENTARY MATERIALS AND METHODS

### Microscopy

All sptPALM measurements were performed on a custom-built microscope platform, designed for interdisciplinary research approaches, using different spectromicroscopy techniques. The following description will therefore also include techniques beyond sptPALM. A complete description of this microscope and its application in various research projects can be found in zur Oven-Krockhaus, 2021. References to specific modules and components refer to Supplemental Figure 1.

### Excitation module

Four different continuous wave (CW) lasers are implemented in the main excitation path: 405 nm (iBeam smart 405-S-LP, 100mW, Toptica), 488 nm (Oxxius Laserboxx Diode laser, 100 mW, OXX-488-100, Laser 2000), 561 nm (Vortran Stradus DPSS laser, 50 mW, VOR-561-050) and 642 nm (Oxxius Laserboxx Diode laser, 130 mW, OXX-642-130). They cover the most common range of available and utilized fluorophores. For techniques requiring a pulsed laser source, a supercontinuum fiber laser (SuperK Extreme EXB-4, with multi-line tunable filter SuperK SELECT UV/VIS-nIR, both NKT Photonics) is also implemented. Excitation intensities are mainly controlled by an acousto-optical tuneable filter (AOTF, Polychromatic Modulator 450 - 700 nm, AOTFnC-VIS-TN, with 4-channel RF driver AA.MPDS4C, both AA OPTO-ELECTRONIC) allowing for the free selection of up to eight laser lines from 400 to 675 nm, which can be used separately as well as in any combination. The CW lasers pass respective clean-up filters (Laser Clean-Up Filter ZET 405/20, F49-405; Laser Clean-up Filter ZET 488/10, F49-488; 560/14 BrightLine HC, F39-561; Laser Clean-up Filter ZET 640/10, F49-643; all AHF analysentechnik AG) and are combined into one beam path using beam splitters (BS1: Beamsplitter HC BS 555, F38-555; BS2: Laser Beamsplitter zt 488 RDC, F43-088; BS3: Laser Beamsplitter H 405 LPXR, F48-403; all AHF analysentechnik AG). Magnetic mirror mounts (M1-3: MDI-HS-3030-M6 on RD-MP magnetic plates, all Radiant Dyes) are used throughout the setup to change excitation or emission paths.

### Coupling module

After passing optional neutral density filters, all lasers are coupled into a single mode fiber (Polarization-Maintaining FC/PC Fiber Optic Patch Cable PM-S405-XP, P1-405BPM-FC-2, Thorlabs) with aspheric lens L1 (350 - 700 nm, f = 11.0 mm, NA = 0.25 Aspheric Lens, C220TMD-A, Thorlabs) for uniform, spatially cleaned-up Gaussian laser profiles. Fiber de-coupling is done with achromatic lens L2 (Ø1” Achromatic Doublet, ARC: 400-700 nm, f=30 mm, AC254-030-A-ML, Thorlabs) collimating the excitation beam to a diameter of 10 mm. Afterwards, additional optical elements like linear polarizers (Mounted GT Polarizer, 10 mm x 10 mm, 350 - 700 nm AR Coating, GTH10M-A, Thorlabs) and/or retardation plates (Ø1/2” Mounted Achromatic Half-Wave Plate, Ø1” Mount, 400 - 800 nm, AHWP05M-600, Thorlabs) can be introduced to control the polarization for respective experiments (e.g. anisotropy measurements). An xyz-adjustable lens L3 (Ø1” UVFS Plano-Convex Lens, f = 150.0 mm, ARC: 350 - 700 nm, LA4874-A-ML, Thorlabs) can be pushed into the beam path, which focuses on the back focal plane of the microscope objective, thus allowing switching between either confocal, or epi-fluorescence/TIRFM/VAEM illumination. For VAEM, the angle can be adjusted by lateral translation of this lens. Laser excitation is reflected by the multi-band beam splitter BS4 (TIRF Quad zt405/488/561/640rpc, F73-410, AHF analysentechnik AG), and directed upwards by a 45° mirror.

### Sample stage

The sample stage mainly consists of a custom-manufactured aluminum frame that holds a piezo table (3-Axis Piezo Scanner with Direct Position Measuring, P-527.3CD, controlled by 3-Axis Digital Piezo Nanopositioning Controller, E-710.3CD, both Physik Instrumente (PI)) for sample-scanning in confocal measurements or small adjustments of the sample position in widefield applications. Laser excitation passes the microscope objective (Objective alpha Plan-Fluar 100×/1,49 Oil M27, 421190-9800-000, Carl Zeiss Microscopy) that can be adjusted in z for focusing onto the sample. An aluminum plate is mounted onto the piezo table with a round recess for the objective, allowing to install different kinds of sample holders. Fluorescence from the sample is collected by the same objective and passes the beam splitter BS4, entering the emission path.

### Detection module

Here, light from the main emission beam path can be redirected via removable mirrors M2 and M3 to the individual detectors. For widefield applications, M2 directs the emission to an sCMOS camera (Digital CMOS camera, ORCA-Flash4.0 V2, Hamamatsu Photonics), focused by an achromatic lens (Ø1” Achromatic Doublet, ARC: 400-700 nm, f=100.0 mm, AC254-100-A-ML, Thorlabs). If needed, green/red channel splitting can be done by the alternative setup shown in Supplemental Figure 1, using beam splitters (BS5: Laser Beamsplitter H 560 LPXR superflat, F48-559; BS6: Shortpass beamsplitter HC BS 556 SP, F38-556; both AHF analysentechnik AG) and two lenses L4 (Biconvex lens; N-BK 7; D=25.4; F=100; mounted, G063854000, Qioptiq) to focus the two emission beams on laterally displaced regions on the camera chip.

For spectroscopy using confocal illumination, the next detector is a spectrograph, consisting of a 30 cm focal length imaging monochromator (Imaging Spectrograph, Acton SP-2356, Princeton Instruments) and a thermoelectrical cooled CCD camera (Digital CCD Camera System, PIXIS 100B, Princeton Instruments), with achromatic lens L5 (Achr. VIS ARB2; D=12.5; F=40; mounted, G052010000, Qioptiq) focusing on the entrance slit.

The following detector block features single-photon avalanche diodes (SPADs, PDM, Micro Photon Devices) for their use in confocal techniques like fluorescence lifetime imaging microscopy (FLIM), fluorescence (lifetime) correlation spectroscopy (F(L)CS), antibunching or other experiments with single-molecule detection. Both SPAD and pulsed laser synchronization signals are fed into a time-correlated single photon counting (TCSPC) module (PicoHarp 300, Picoquant GmbH) to measure the time delay between sample excitation and the arrival of an emitted photon at the detector. Insets a-e in Supplemental Figure 1 describe different configurations using different lenses (L6: Ø1” Achromatic Doublet, ARC: 400-700 nm, f=40.0 mm, AC254-040-A, Thorlabs; L7: Achr. VIS ARB2; D=25.4; F=200; mounted, G063237000, Qioptiq; L8: Achr. VIS ARB2; D=25.4; F=400, G322340322, Qioptiq) for specific experimental setups: (a) dual-color FLIM/F(L)CS with beam splitter BS7 (Laser Beamsplitter H 560 LPXR superflat, F48-559, AHF analysentechnik AG); (b) anisotropy measurements with polarizing beam splitter BS8 (20 mm Polarizing Beamsplitter Cube, 420 - 680 nm, PBS201, Thorlabs); (c) low intensity measurements (no optical sectioning to maximize signal intensity); (d) high-contrast imaging (pinhole for optical sectioning); (e) compromise between (c) and (d), the projected Airy disk is about the size of the SPAD’s detector area, effectively excluding most out-of-focus light.

### Hardware control

All lasers are controlled (directly and via the AOTF) by a custom-written software in LabVIEW (LabVIEW 2018, National Instruments). A standard PC gaming controller is used for focusing and (for widefield applications) lateral adjustments of the sample position, also written in LabVIEW. Widefield sCMOS camera data is recorded with the HoKaWo software (HoKaWo 3.0, Hamamatsu Photonics), while TCSPC data is acquired with SymPhoTime 64 (SPT64-1+2, Picoquant GmbH).

